# Climate-driven niche tracking and genomic resilience shape future distribution of a widespread agricultural weed

**DOI:** 10.64898/2026.08.07.743277

**Authors:** Célia Neto, Quanjing Zheng, Paul Neve

## Abstract

Understanding how agriculturally important species respond to environmental change is critical for maintaining productivity, mitigating agroecosystem threats, and sustaining resilience. While crops have traditionally been the focus in agroecosystems, agricultural weeds are integral components that often face even stronger selective pressures, making them powerful models for investigating ecological and evolutionary responses to climatic and human-mediated challenges. Insights from how weeds adapt rapidly under these pressures can inform strategies to improve agricultural outcomes, since both pests and crops evolve under the same multivariate selective pressures. Here, we integrate two centuries of distribution records with whole-genome sequencing from natural populations of the most damaging weed in Europe – *Alopecurus myosuroides* (blackgrass) – to examine its ecological and evolutionary responses in agroecosystems. Blackgrass largely maintained its historical climatic niche, expanding its range primarily by tracking environments analogous to those it historically occupied. Genome-wide analyses revealed a polygenic basis of environmental responses, with most loci linked to single environmental variables and a subset showing limited environmental pleiotropy, indicating modular adaptation to the complex selective pressures of managed agricultural landscapes. Coupling these genomic-environment relationships with projected climate change and genomic offset analyses indicated that most blackgrass populations will remain well aligned with future conditions. Our findings show that ecological niche tracking and polygenic adaptation allow agricultural weeds like blackgrass to persist under rapid environmental change, offering insights relevant not only for weed management but also for designing resilient cropping systems under future climates.

## Introduction

Globally, climate and land-use change are driving rapid shifts in plant species distributions (Guo et al., 2018; Parmesan & Yohe, 2003; Rubenstein et al., 2023; Thuiller, 2007). In agroecosystems, these dynamics are often intensified as abiotic factors (climatic and edaphic) interact with biotic agents and agricultural management to generate novel and intense selective pressures. Agricultural weeds evolve exceptionally rapidly under recurrent human-mediated selection and occupy landscapes where climate, soils, and cropping systems jointly shape adaptation, making them powerful models for understanding the genomic eco-evolutionary responses that enable plants to persist and spread in rapidly changing environments (Kreiner et al., 2022; Mahaut et al., 2020; Baucom, 2019, 2016; Peters et al., 2014; Neve et al., 2009).

Historical and contemporary changes in species distributions can arise through niche tracking, whereby species distributions track shifting environmental conditions, or by niche evolution, whereby species evolve changes in environmental tolerances (Waldvogel et al., 2020). Because these responses depend on combinations of dispersal capacity, physiological limits, and genetic variation, they have distinct implications for forecasting species responses under accelerating global change (Guo et al., 2018; Parmesan & Yohe, 2003; Rubenstein et al., 2023; Waldvogel et al., 2020). In agricultural weeds, resolving the relative importance of ecological, evolutionary, and genetic responses to environmental change is important for anticipating their future ecological and agricultural impacts.

Although ecological niche models (ENMs) are powerful tools for projecting species distributions (Alvarado-Serrano & Knowles, 2014; Peterson, 2006), they typically assume that species-environment relationships remain constant through time. In addition, the genetic variation and adaptive potential within and among populations that may enable persistence under novel environmental combinations or strong anthropogenic selection are often overlooked (Aguirre-Liguori et al., 2021; Broennimann et al., 2007; Guisan et al., 2014; Guisan & Thuiller, 2005; Soberón & Nakamura, 2009; Thuiller et al., 2008; Waldvogel et al., 2020). Landscape genomics provides a complementary framework by linking allele frequency variation to environmental gradients, which can then be used to quantify genomic offset – the genomic change required to maintain adaptation under future environmental conditions (Aguirre-Liguori et al., 2021; Capblancq et al., 2020; Waldvogel et al., 2020). Integrating ENMs and genomic offset provides a temporal framework for testing whether historical range shifts primarily reflect niche tracking or niche evolution, and whether contemporary populations harbour sufficient adaptive variation to persist under future climates.

Adaptive responses to environmental change also depend on the genomic architecture of adaptation, including the number, effect sizes and pleiotropy of loci contributing to adaptation (Bomblies & Peichel, 2022; Connallon & Hodgins, 2021; Höllinger et al., 2019; Orr, 2005). While rapid adaptation can be underpinned by both polygenic and monogenic architectures (Gomulkiewicz & Holt, 1995), the former is expected to be favoured under rapid environmental change with milder shifts (Bellagio & Exposito-Alonso, 2025; Kardos & Luikart, 2021). Of relevance is to consider whether adaptive loci influence multiple environmental axes – environmental pleiotropy – or show more modular genetic architectures, whereby adaptive loci respond independently to different environmental factors. This will dictate how responses to one component of environmental change constrain or enable adaptation to others (Lotterhos et al., 2018).

We focus on *Alopecurus myosuroides* (blackgrass), an annual grass native to western Asia and the eastern Mediterranean. Likely introduced to Europe during the last 10 000 years, blackgrass was already recognized as a “very troublesome weed among wheat” by the early 19^th^ century (Sinclair, 1838). Today, blackgrass is widely distributed across Europe and is a major yield-limiting weed of autumn-sown cereal crops (Ahmad et al., 2021; Cai et al., 2023; Hicks et al., 2018; Moss, 2017). Its broad geographic and climatic distribution makes it an excellent model for studying local adaptation, niche evolution and spatiotemporal range dynamics.

We address two main questions: (i) how has blackgrass expanded its range over the last two centuries? and (ii) will it persist under future climate change? To answer these, we reconstruct historical range dynamics, quantify past and current environmental suitability, and test for shifts in the species’ ecological niche. We then assess whether range expansion reflects niche evolution or niche tracking, identify the environmental drivers, and use whole-genome sequencing data from 64 blackgrass populations to estimate genetic vulnerability under projected future climates. For blackgrass, these findings will help to determine whether it will remain a persistent threat under future climates. More broadly, this work contributes to emerging efforts to implement ecological landscape genomics to forecast evolutionary responses in rapidly changing environments, including agroecosystems.

## Material and Methods

All analyses, including data collection, niche modelling, statistical analyses, and genomic offset, were conducted in R v.4.4.1 (R Core Team, 2021) with custom scripts, available at https://github.com/celianeto/blackgrass_climate/. The bioinformatic pipeline for processing of genomic data is available in the same GitHub repository with detailed parameters and flags used. Supplemental Information contains description of further analyses.

### Species occurrence data

Data for blackgrass presence in Europe and the Middle East (latitude 30° to 70° North and longitude −10° to 50° West, 1900 to 2021) was retrieved from the Global Biodiversity Information Facility (GBIF, http://www.gbif.org/) (GBIF.Org, 2022a, 2022b), with the parameters *Has coordinate: true*, *Has geospatial issue: false*, *Scientific name: Alopecurus myosuroides Huds.*, and *Year: Between start of 1900 and end of 2021*. Only records where both location and date of collection were available were kept. The final data set was split into ‘past’ (collection date 1900 – 1921) and ‘present’ (2000 – 2021). These two time periods were chosen to capture contrasting climatic and land-use conditions before and after the major agricultural intensification of the 1960’s, and to match available environmental data (see below).

To account for overrepresentation, potential taxonomic misidentifications or erroneous coordinates, we grouped blackgrass occurrences into 0.5° x 0.5° cells. For each cell, independently from how many records we had initially, we retained a single record, chosen randomly. After applying spatial thinning to reduce uneven sampling intensity and minimise the influence of potential misidentifications, the dataset retained 44 historical and 443 contemporary occurrences. These thinned datasets were used for all downstream analyses.

To test whether the increase in blackgrass occurrences could be driven by increased recording effort, we conducted two complementary analyses using *Poaceae* records as a proxy for sampling effort (as blackgrass belongs to the *Poaceae* family).

First, we modelled thinned blackgrass presence/absence across all grid cells as a function of period (past or present) using a logistic regression, with log-transformed *Poaceae* record count per cell as a covariate for effort intensity. The effect of period was assessed by a likelihood ratio test against a model containing only the sampling effort covariate. Second, we performed a rarefaction analysis by repeatedly subsampling present-day occurrences to match past sample size and compared the number of occupied grid cells across 1000 replicates. An empirical p-value was calculated as the proportion of replicates occupying fewer or equal cells than the past median.

### Environmental data

Environmental data consist of climatic, edaphic and land-use data for the geographic region considered, and were all obtained at or transformed to 30 arc-minutes spatial resolution (equivalent to 0.5° x 0.5°).

To assess temporal changes, we compared two 20-year time windows representing past and present climatic conditions. The Past period (1900-1921) was selected as the earliest interval for which high-resolution bioclimatic variables are available (from BioClim), while the Present period (2000-2021) represents contemporary conditions with comparable data coverage. Using 20-year intervals helps smooth interannual climatic variability and provides a representative average for each period. The two windows are separated by approximately 80 years to maximize climatic contrast and to allow sufficient time for potential eco-evolutionary responses. These time windows are pre- and post-agricultural intensification and widespread herbicide use that drastically altered the evolutionary trajectories of weed species.

Climatic data encompass the 19 BioClim variables calculated based on monthly maximum and minimum temperatures and precipitation, from CRU TS v4.07 (Harris et al., 2020), using the *biovars()* function in the dismo package (Hijmans et al., 2010) in R. Edaphic data were downloaded with the package geodata (Hijmans et al., 2025) and are derived from the SoilGRIDS database for a soil depth of 30-60 cm collected in 2003 (Poggio et al., 2021). The data include values for total nitrogen (g/kg), percentage of clay, silt and sand in fine earth, pH and bulk density of the fine earth fraction (kg/dm^3^). A per area estimate of cropland use was derived from the HYDE 3.5 database (Klein Goldewijk et al., 2026), given as km² of cropland per grid cell. To account for uneven data coverage over the past and present periods, we averaged the available decadal layers for the past period (1900 and 1910), and the start and end layers of the present period (2000 and 2021).

For each record, environmental data were averaged across the four adjacent cells surrounding the collection site, and across months and years for the corresponding time period. Correlation matrices were calculated for all 26 environmental variables using Spearman correlations (function *cor()*; Supplemental Figure 1), and their similarity was assessed using a Mantel test (function *mantel()* from vegan package) (Oksanen et al., 2001). Since the two were statistically similar (Mantel test, r = 0.88, p-value = 0.001), present-day data were used for variable selection. To minimise collinearity, variables were selected using the variance inflation factor (VIF) method, from the usdm package (Naimi et al., 2014). Using the function *vifcor()*, with a Spearman correlation threshold of R = 0.5, only variables below this threshold were retained. This conservative threshold was chosen to ensure predictor independence, as downstream environmental pleiotropy analyses are particularly susceptible to inflation by collinear predictors.

After accounting for collinearity, six environmental variables were retained. These were the BioClim variables bio4 (temperature seasonality), bio8 (mean temperature of the wettest quarter), bio13 (precipitation of the wettest month), bio15 (precipitation seasonality), the edaphic variable soil silt percentage, and cropland use.

### Centroid, Overlap, Unfilling, and Expansion framework

To assess blackgrass responses to environmental change, we employed the Centroid shift, Overlap, Unfilling, and Expansion (COUE) framework (Broennimann et al., 2014). This framework provides quantitative metrics of niche stability, expansion, and unfilling, to describe species spatio-temporal responses to environmental change.

Niches were estimated by applying a kernel smoother function to the density of past and present species occurrences in gridded environmental space along the first two PCA axes, using the function *ecospat.grid.clim.dyn()* from the ecospat package (Broennimann et al., 2014). We decomposed niche changes into niche stability, niche expansion, and niche unfilling. Niche stability is defined as the proportion of the current niche that overlaps with the past niche in environmental space, while niche expansion (1-stability) is the proportion of the current niche occupying novel environments. These two metrics provide insight into the potential evolutionary responses underlying niche change: high niche stability suggests niche tracking, whereas high niche expansion is consistent with niche evolution. Finally, niche unfilling quantifies environmental space that is occupied in the past range but not in the current range. It was calculated as the proportion of past occurrences located in climate conditions that remain available but are unoccupied in the current range. Overall niche overlap was measured using Schoener’s D (Schoener, 1970), which ranges from 0 (no overlap) to 1 (complete overlap) using the *ecospat.niche.overlap()* function from ecospat. We then used Schoener’s D to test for niche conservatism during range expansion using niche equivalency and niche similarity tests (Broennimann et al., 2012; Warren et al., 2008).

The niche equivalency test (function *ecospat.niche.equivalency.test()* from ecospat) randomly reallocates occurrences between the two niches to test whether they are identical, whereas the niche similarity test (function *ecospat.niche.similarity.test()* from ecospat) evaluates whether the two niches are more similar to each other than to other niches selected at random from the study area. Both tests were repeated 1000 times to generate a null distribution and to confidently reject/accept the hypothesis of niche conservatism.

To assess whether niche estimates were sensitive to the imbalance in sample size (past = 44, present = 443), we performed two complementary analyses. First, we randomly downsampled the present-day records to match the number of past records and recalculated all niche statistics, repeating this procedure 100 times. We then calculated empirical p-values as the proportion of iterations yielding values at least as extreme as those observed on the full data set. Second, to test whether past records adequately captured the full breadth of the historical niche, we constructed a nested dataset combining both past and present occurrences and recomputed niche dynamics and overlap statistics against the present-day environment.

### Maximum Entropy modelling

MaxEnt (Maximum Entropy modelling) estimates habitat suitability from presence-only occurrence data by contrasting environmental conditions at occurrence localities with those available across the study area (Phillips et al., 2006). We used MaxEnt to address two questions: how has the predicted distribution of blackgrass changed between the two study periods, and what was the relative contribution of climate versus agricultural land-use explaining its current distribution.

Using the maxnet R package (Phillips, 2022) and the function *maxnet()*, we initially built three models: (i) the Past model – past occurrences mapped onto past environmental variables, (ii) the Present model – present-day occurrences on present-day environmental variables, and (iii) the Projected model – past occurrences projected on present-day environmental variables. Occurrence data for both time periods were taken from GBIF and BioClim variables from CRU (see above). We then created a fourth model, the Future model, in which present-day occurrences were projected under future climate scenarios. Future models were built for 2061-2080, using combinations of three global climate models (CNRM-CM6-1, MPI-ESM1-2-LR, and UKESM1-0-LL) and three shared socio-economic pathways (SSPs 126, 245, and 585, representing “sustainability”, “business as usual”, and “worst case”, respectively). Future climate projections were obtained using the function *cmipC_world()* from the geodata package (Hijmans et al., 2025) in R, and the resulting rasters were aggregated to 0.5° resolution to match past- and present-day climate layers. Background points (n = 5000) were randomly sampled from the full study extent (function *spatSample()*, terra package) (Hijmans et al., 2026). Models were fitted using linear, quadratic, and hinge response functions (option classes = “lqh”). Model performance was assessed using 100 bootstrap replicates with an 80/20 train/test split, and reported as the mean AUC ± 95% confidence interval (function *roc(),* pROC package) (Robin et al., 2011).

We assessed the relative contribution of climate and agricultural land-use to blackgrass distribution in the past and present by fitting a combined MaxEnt model (climate + agriculture) at each time period. Within each combined model, variable importance was quantified for each predictor by randomly permuting its values across observations and regenerating predictions from the permuted data; the resulting decrease in AUC relative to the unpermuted model was recorded as that variable’s importance.

### Genomic data

Whole-genome sequence data were generated using a pool-seq approach for 64 blackgrass populations from across the species range (Supplemental Table 1). For each population, we sampled a 2 cm leaf section from 25 individuals, pooled these leaves and extracted pooled DNA using the DNeasy Plant Mini Kit (Qiagen), following manufacturer’s protocol. This pooling strategy represents a compromise between genome size (3.6 Gb), sequencing depth, population number, and cost, prioritising broad geographic and climatic coverage over deeper per-population sampling. These 64 pools (1600 individuals in total) were whole-genome sequenced, using Illumina 150 bp pair-ended short-reads.

Raw reads were trimmed using Trimmomatic (“LEADING:5” “TRAILING:5” “SLIDINGWINDOW:4:20” “MINLEN:50”) (Bolger et al., 2014). Trimmed reads were mapped to the *Alopecurus myosuroides* reference genome (NCBI version: JAPCYS010000000) (Cai et al., 2023), using the maximal exact matches algorithm implemented in bwa (Li, 2013). Resulting sam files were further manipulated using samtools (Danecek et al., 2021), to create sorted bam files (using *view* and *sort* with -q 20 for quality filtering). Duplicates were marked and removed using picard (http://broadinstitute.github.io/picard). Per population bam files were combined into a single mpileup file with samtools *mpileup*. Sync files were created with popoolation2 with mpileup2sync.jar (Kofler et al., 2011) and converted into genobaypass files with the functions *popsync2pooldata()* and *pooldata2genobaypass()* in the R package poolfstat (Hivert et al., 2018). SNPs were filtered by requiring a minimum read count of five per population (*min.rc = 5*) and a minimum minor allele frequency of 0.05 (*min.maf = 0.05*). The resulting files were then used as input in BayPass (Gautier, 2015) for genetic mapping. Unless otherwise stated, default parameters were used. In total, 103 888 478 SNPs were retained, with a mean coverage of 14.5x per pool (minimum across pools = 10.16, maximum = 18.78; Supplemental Table 1).

### Genome-environment association

We performed genome-environment association (GEA) analyses to identify genetic variants associated with the six environmental variables contributing to the niche of blackgrass. GEA was initially conducted using the standard covariate model in BayPass (Gautier, 2015), which implements a Bayesian hierarchical model to account for covariance among populations due to shared evolutionary history, improving the detection of true environment-allele associations. We used the present-day values for the environmental variables at the geographical origin of each of the 64 blackgrass populations as phenotypes. Following association analyses, p-values were corrected for linkage disequilibrium using the local score approach to identify genomic regions significantly associated with each environmental variable (Bonhomme et al., 2019). We also calculated BayPass XtX values for each SNP. XtX quantifies genetic differentiation among populations and can be used to identify loci showing high differentiation consistent with local adaptation (Gautier, 2015).

To assess the robustness of the genetic architecture inferred by BayPass, we employed two additional GEA methods, LFMM and RDA (Supplemental Information). All three approaches supported a strongly polygenic architecture of climate adaptation, although overlap among candidate SNPs was limited. This reflects fundamental differences between methodologies, including how each approach models pool-seq sampling variance, corrects for population structure, and handles linkage disequilibrium (further discussion in Supplemental Information). As BayPass was specifically designed for pool-seq data, we used its results for all downstream analyses.

We identified all genes in each of the significant regions identified (n = 3523, across all six environmental variables), including genes in the regions 10 kb upstream and downstream. These were treated as candidate genes. For each of them, we looked for a corresponding ortholog in *Arabidopsis thaliana* (reference TAIR10) using OrthoFinder (Emms & Kelly, 2019), and assessed gene ontology enrichment in this set of genes with the Classification SuperViewer Tool w/ Bootstrap (Provart & Zhu, 2003) with 100 bootstraps.

When the same gene showed up as candidate in association with different environmental variables, we considered it as environmentally pleiotropic (Lotterhos et al., 2018). Genes intersection between variables were assessed with upset plots (function *upset()* from package UpSetR) (Conway et al., 2017) in R. To assess whether the number of overlapping genes between environmental variables was greater than expected by chance, we generated a null distribution based on 1000 random genome draws. For each environmental variable, we randomly sampled *s* SNPs from the genome, where *s* is the number of significant regions identified for that variable in our GEA analysis. Each sampled SNP was annotated to its nearest gene using a 10kb-flanking window (the same procedure used for the observed associations). We then quantified gene overlap among variables for all random draws and compared these values to the observed overlap to compute empirical p-values. Genetic correlation between environmental variables at the SNP-level were based on effect size (beta) estimated by BayPass.

### Genomic offset

We quantified potential maladaptation to future environmental change using the geometric genomic offset (Gain et al., 2023) and risk of non-adaptedness (RONA) (Rellstab et al., 2016) approaches, using the function *compute_genetic_offset()* from BayPass (Gautier, 2015). For each genomic region associated with an environmental variable, we selected the SNP with the lowest p-value and used its estimated effect sizes for all mapped environmental variables across populations to create a genomic matrix. Then, we calculated an environmental matrix, as the mean difference between environmental variables in present and projected future climates. Geometric genomic offset was calculated as the mean, across all SNPs, of the squared environmental difference between present and future conditions weighted by each SNP’s effect sizes (Gain et al., 2023), while RONA was calculated as the mean, across all SNPs, of the absolute value of this same weighted difference (Rellstab et al., 2016; Sang et al., 2022). High genomic offset values indicate that substantial shifts would be necessary to maintain local adaptation under projected future climates, while low values indicate that existing genetic composition is already well matched to future environmental conditions (Aguirre-Liguori et al., 2021). We built geometric genomic offset models using the future climate scenarios outlined above.

To assess the sensitivity of genomic offset estimates to candidate SNP selection, we recomputed genomic offset using three candidate SNP sets of increasing stringency: all candidate loci, the top 10% by XtX, and the top 1% by XtX.

## Results

### Blackgrass distribution tracked shifting environmental conditions

The earliest recorded occurrence of blackgrass in Europe was from 1750 in Germany, but it remained relatively rare for nearly a century, with its frequency increasing in herbarium collections from the mid 1800’s onwards (GBIF). More recently, at the end of the 20^th^ century, a substantial rise in occurrences has been documented, based on both herbarium records and citizen science reports (Figure 1). This increase in recorded occurrences reflects a genuine increase in distribution rather than simply increasing recording efforts, as shown by a significant effect of period of collection after controlling for effort intensity (logistic regression, likelihood ratio test, p-value < 2.2×10^-16^), and by a rarefaction analysis (empirical p < 0.001).

**Figure 1.**
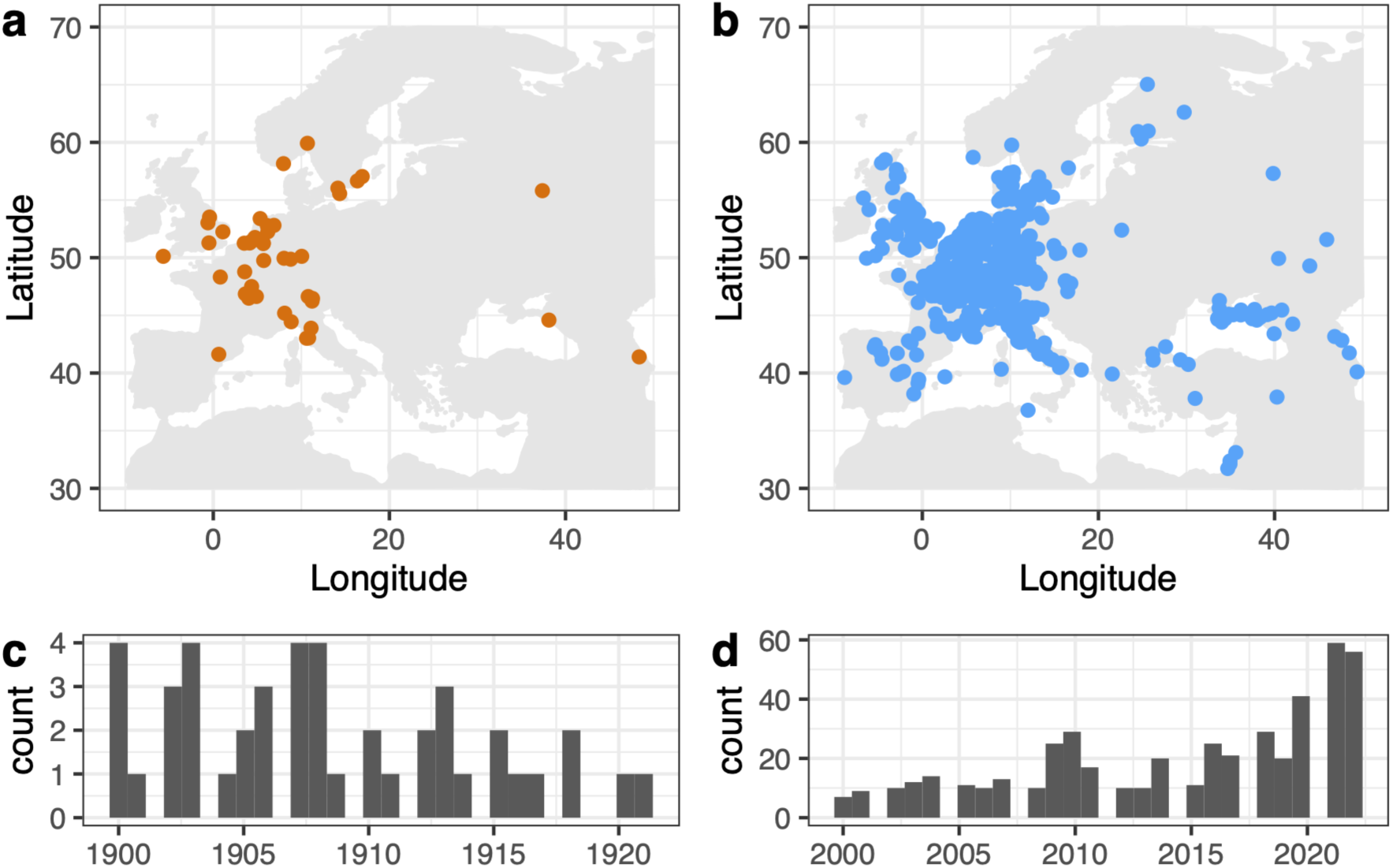
Blackgrass distribution across Europe in the last two centuries. **a.** and b. Geographical distribution of blackgrass in the past (1900 to 1921, n = 44) and in the present (2000 to 2021, n = 443), respectively. c. and d. Temporal distribution of blackgrass records across Europe in the past and present time periods, respectively. All data were retrieved from GBIF.

Ecological niche comparisons between the past (1900-1921) and present (2000-2021) using the COUE framework and six non-collinear environmental variables (Supplemental Figure 1) revealed high niche stability (94%) and limited niche evolution (6%) (Figure 2). This was supported by strong niche overlap (Schoener’s D = 0.66, similarity test p-value = 0.003, equivalency test p-value = 0.406; Figure 2a), indicating that contemporary occurrences largely occupy environmental conditions consistent with those inferred from historical records. Together, these results suggest that blackgrass has primarily tracked suitable environmental conditions during the last century, with limited evidence for niche evolution.

**Figure 2.**
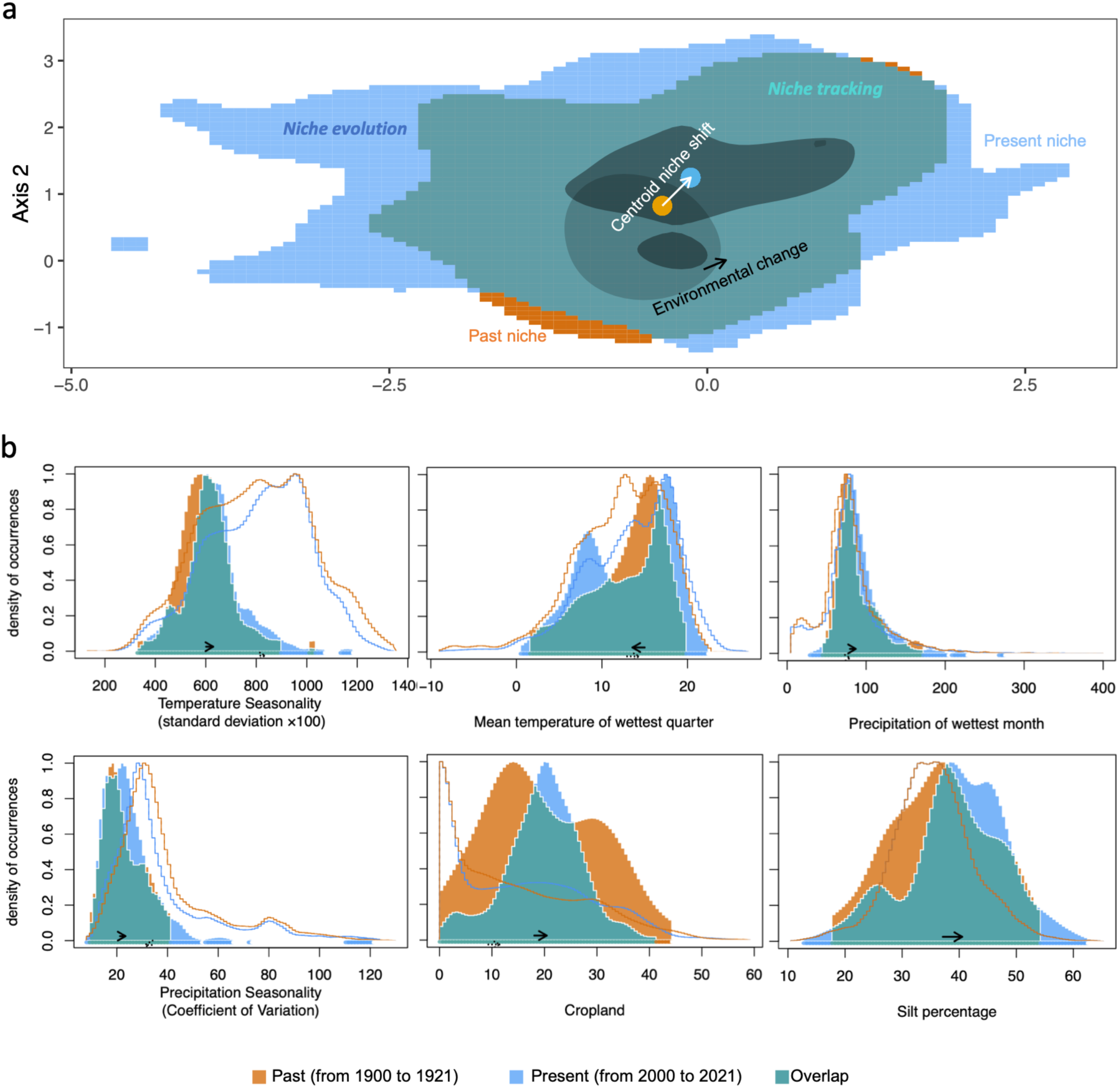
Blackgrass niche dynamics over the last two centuries. a. Past (orange) and present (blue) environmental niches of blackgrass in environmental space. The overlap between past and present niche space (niche stability) is shown in green. The black arrow indicates the shift in the environmental centroid shift between past and present climates, while the white arrow indicates the shift in blackgrass niche centroid from the past (orange dot) to the present (blue dot). Dark and light shading within the area of niche stability indicates areas with the highest density of past and present blackgrass occurrences, respectively. b. Distributions of environmental variable values between the past (orange) and present (blue). The x-axis shows the values of each environmental variable and y-axis the density of occurrences for each environmental value in the past and present. Solid lines indicate the extent of environmental conditions occupied in the past (orange) and present (blue). Soil silt percentage extent show overlapping distributions, as it was derived from a single time point (2003). Throughout the figure, green represents overlap between past and present niches, orange represents past niche only, and blue indicates the present niche only.

Downsampling present-day occurrences to match the historical sample size (n = 44) had no significant effect on niche estimates (Supplemental Table 2), indicating that these are robust to differences in sample size. Similarly, recomputing niche dynamics using a nested data set (past + present) produced estimates (stability = 94%, expansion = 6%, unfilling = 0%, Schoener’s D = 0.71) that closely matched the time-binned analysis, suggesting that the historical records adequately captured the breadth of the historical niche.

Over time, blackgrass occurrences shifted into areas with warmer wet seasons, increased precipitation, and more stable precipitation regimes (Figure 2b). They also shifted towards areas with reduced temperature seasonality, counter to the general climate trajectory. Blackgrass also expanded into cropland-dominated areas with higher soil silt content, suggesting that both climatic and agricultural factors have contributed to its current distribution.

### Environmental change has reshaped blackgrass distribution in Europe

To assess how environmental change reshaped blackgrass distribution across Europe, we constructed Past, Present and Projected models (Table 1, Figure 3) using MaxEnt. First, we quantified the relative contribution of climate vs agricultural land-use to blackgrass distribution changes across time. Climate variables showed consistently higher permutation importance than cropland in both periods (past: climate = 0.377 vs. cropland =0.019; present: climate = 0.242 vs. cropland = 0.113), indicating that climate remained the dominant predictor of blackgrass distribution throughout. However, cropland’s importance increased substantially over the 20^th^ century. Nonetheless, because climate and agricultural intensification have co-varied over this period, their effects cannot be fully disentangled.

**Figure 3.**
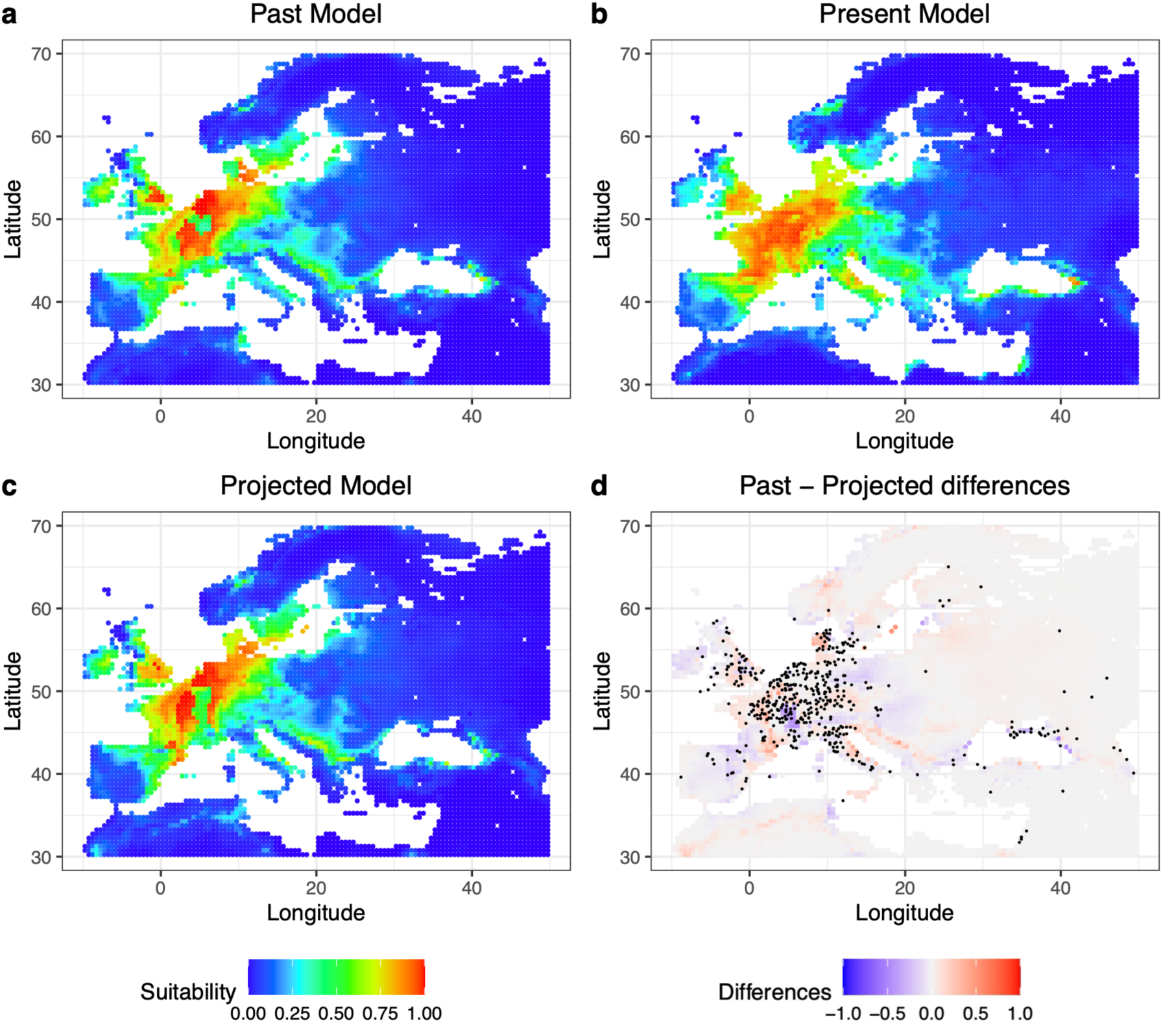
Blackgrass’s predicted suitable habitats across Europe in the last 200 years. Suitable habitats predicted by MaxEnt under a. the Past model (past occurrences on to past environmental data), b. the Present model (present-day occurrences on present-day climate), and c. the projected model (past occurrences on to present-day environment). Colour refers to predicted suitability following the legend. d. Differences in habitat suitability between the Past and the Projected models. Colour refers to differences in predicted suitability between the two models considered. A positive value represents higher suitability predicted under the Projected model.

**Table 1.**
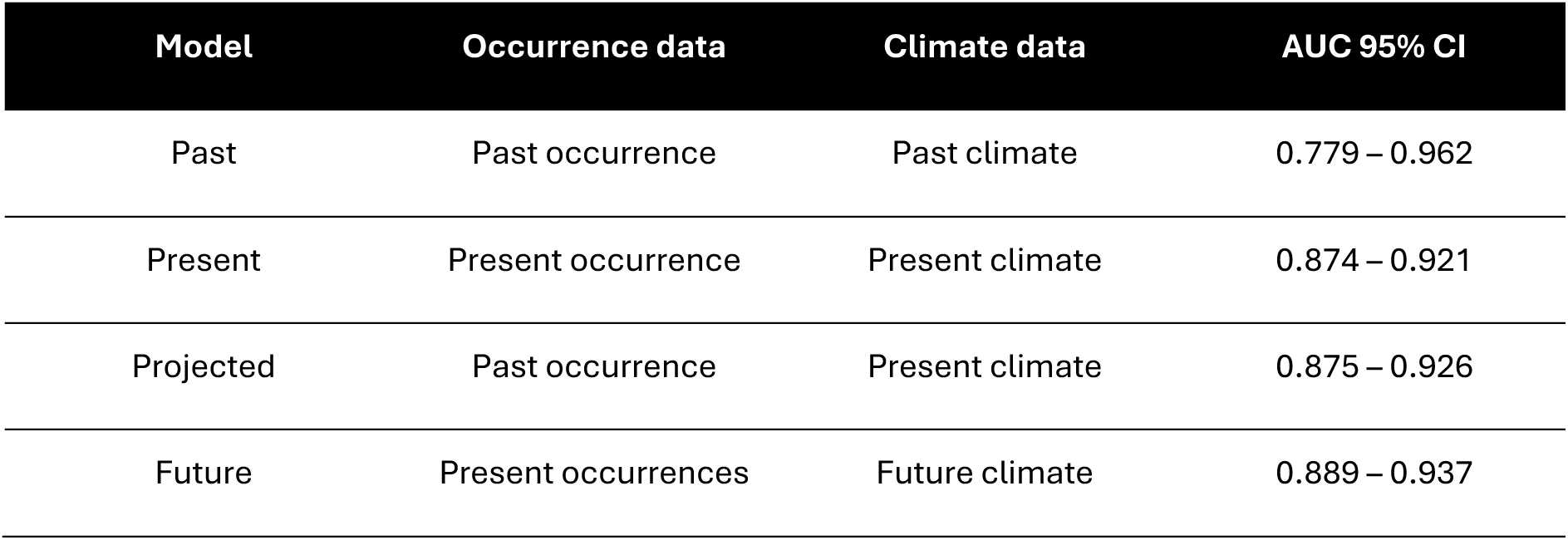
MaxEnt models. Description includes model name, occurrence and climate data used in each of the models, and their corresponding 95% confidence interval over 100 bootstraps for AUC calculation.

| Model | Occurrence data | Climate data | AUC 95% CI |
| --- | --- | --- | --- |
| Past | Past occurrence | Past climate | 0.779 – 0.962 |
| Present | Present occurrence | Present climate | 0.874 – 0.921 |
| Projected | Past occurrence | Present climate | 0.875 – 0.926 |
| Future | Present occurrences | Future climate | 0.889 – 0.937 |

Comparing the Past and Present models identified areas where suitability changed across time (Supplemental Figure 2). Suitability increased (>0.05) across 19% of the study area, mainly around the Adriatic Sea, northern and western Turkey, the Iberian Peninsula, southern and western France, southwest England, and Austria, the Czech Republic and southeast Germany, while it decreased (<-0.05) across 11% of the area, including Ireland, the Baltic region and the Netherlands.

To further distinguish niche tracking from niche evolution, we compared the Past and Projected models (Figure 3d). If blackgrass were simply tracking environmental changes, suitability under the Past and Projected models would be equivalent. A positive change in suitability (i.e., higher under the Projected model) instead identifies regions where environmental change over the last 200 years has created conditions allowing blackgrass to expand (red areas in Figure 3d), while a negative change in suitability identifies regions where environmental change would be expected to render conditions unsuitable (higher suitability in the Past, blue areas in Figure 3d), assuming no change in the species’ environmental relationship. Overlaying current occurrences onto this comparison, however, shows that blackgrass occurs in several of these predicted unsuitable regions, such as the north coast of the Black Sea, the northeast Iberian Peninsula and southwest France, Scotland, Switzerland, Slovakia, Ukraine, and Poland. Its persistence in habitats predicted to be unsuitable based on its past environmental relationships indicates a shift in that relationship, consistent with minimal niche evolution (6% as estimated above) and potential adaptation to novel environmental conditions.

Finally, we used present-day climate-occurrence associations to model blackgrass’s potential distribution under future climate projections – the Future model (2061-2080; Table 1, Supplemental Figure 3). Comparing the Present and Future models (Figure 4) showed that most of Europe is likely to remain suitable for blackgrass, with modest regional shifts: 4% of the area is predicted to gain suitability (>0.05), mainly north of the Alps (Switzerland, Austria, Czech Republic, southeast Germany), in Poland and Belarus, and in southeast England, while 18% is predicted to lose suitability (<-0.05). Projections were consistent across climate models and emission scenarios (Supplemental Figure 4), indicating that this pattern of modest regional shift is robust to climate uncertainty. Whether populations can actually keep pace with these projected changes, however, depends on their capacity for adaptive evolution.

**Figure 4.**
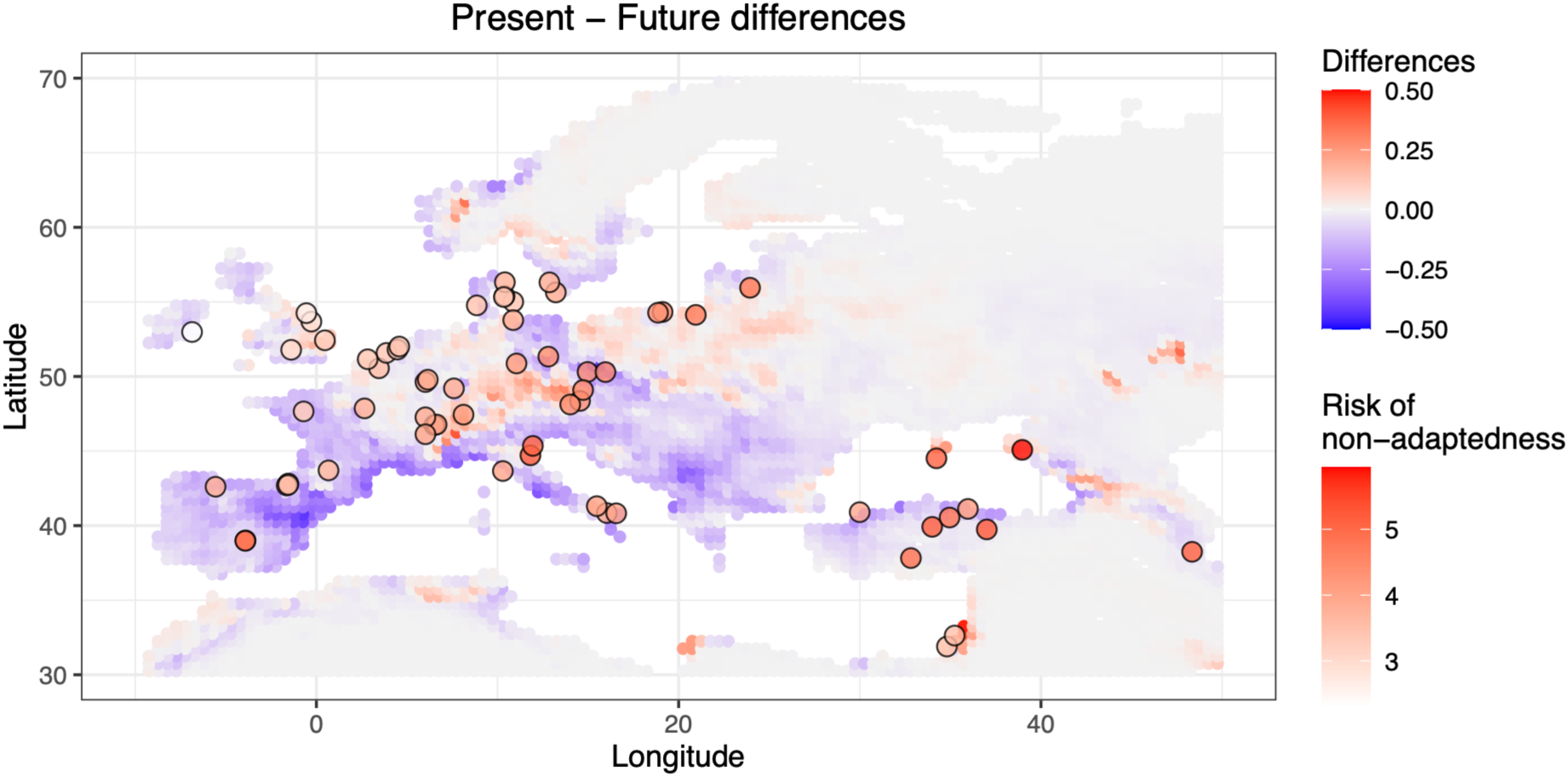
Future maladaptation risk across blackgrass’s range. Map showing predicted suitability differences between Present and Future models. Areas coloured in shades of red show higher suitability under the MaxEnt Future model than under the Present, while blue regions show the opposite. Overlapping, represented by the large dots, are the collection locations of the populations sequenced in this study. These dots are coloured according to the estimated genomic offset value – here as risk of non-adaptedness (RONA). Values correspond to predictions using the MPI-ESM1-2-LR model, for ssp 245 (“business as usual”), between 2061 and 2080. Higher risk of non-adaptedness (more red) indicates populations at higher risk under future climate change, whereas whiter populations were predicted to be less genetically vulnerable.

### Polygenic environmental adaptation supports blackgrass resilience to future climates

Because the previous correlative models cannot capture local maladaptation or limits to adaptive capacity, we next performed a genomic offset analysis, quantifying the predicted mismatch between current genomic composition and that required under future conditions. First, we identified genomic regions associated with climate variation. Using genome-environment association (GEA) analyses across 64 natural populations, spanning the species’ environmental range in Europe (Supplemental Figure 5), we identified a total of 3523 genomic regions associated with environmental predictors (from 261 loci associated with cropland to 1511 genomic regions associated with temperature seasonality; Supplemental Figure 6, Supplemental Table 3). These results indicate a polygenic architecture for climate adaptation. A high number of climate-associated loci was consistently found when GEA analyses were conducted using complementary methods, including LFMM and RDA (Supplemental Information), confirming the polygenicity of climate adaptation.

These candidate climate-associated genomic regions exhibited significantly greater XtX values than genome-wide expectations (one-sided Kolmogorov-Smirnov test, D^+^ = 0.55, p-value < 2.2×10^-16^), consistent with positive selection and local adaptation (Supplemental Figure 7). Within these regions (±10kb), we annotated 1019 genes enriched for biological processes associated with abiotic and biotic stress responses, energy metabolism, development, light and signal-response pathways, and carbohydrate metabolism (Supplemental Figure 8). These pathways suggest that traits relating to growth, phenology, photoperiod response, and biotic interactions may have contributed to the historical range expansion of blackgrass and adaptation to novel and diverse environments.

While most candidate loci were associated with a single environmental variable, 8% showed associations with multiple variables (Supplemental Figure 9). Null-model randomizations revealed significant enrichment (empirical p-value < 0.001) of some multi-variable associations, including temperature seasonality with soil silt percentage, and precipitation seasonality with precipitation of the wettest month (Supplemental Figure 10). One gene (ALOMY6G44813), of unknown function, was associated with four variables (temperature and precipitation seasonality, precipitation of the wettest month, and silt percentage in soil). SNP-level patterns showed similar enrichment of multi-variable associations despite weak correlations between those environmental variables (Spearman’s ρ = −0.06 and 0.12, respectively; Supplemental Figures 11 and 1). Conversely, some combinations of variables showed significantly fewer overlaps than expected (Supplemental Figure 10). Overall, these results indicate a mostly variable-specific, polygenic adaptive architecture with limited environmental pleiotropy, allowing largely unconstrained environmental adaptation.

Finally, we estimated genomic offset to quantify the predicted mismatch between current genomic composition and that required to track future climate conditions, using both geometric genomic offset and RONA (risk of non-adaptedness) based on the environment-associated loci identified above (Supplemental Figure 12). Populations in western Europe showed lower offset values, suggesting that their current genetic composition is well aligned with projected future climatic conditions. Higher offset values occurred in parts of the Black Sea region and isolated southern and central European populations, indicating that larger genomic shifts may be required under future climate (Figure 4).

To assess the influence of potential false positives, analyses were repeated using increasingly stringent candidate SNP sets (all loci, top 10% XtX and top 1% XtX). Results were consistent across all thresholds (Supplemental Figure 13), indicating that genomic offset estimates were robust to candidate SNP selection. Similarly, genomic offset values were consistently low across climate models and emission scenarios (Supplemental Figure 14).

## Discussion

Predicting how agricultural weeds (and other pests) respond to environmental change is important for mitigating their impacts on agricultural productivity in future agroecosystems. By integrating historical niche reconstruction with genome-wide analyses across continental environmental gradients, our study provides evidence that blackgrass, a major agricultural weed of cereal crops, expanded its range across Europe over the past two centuries primarily through niche tracking, with limited niche evolution. Genome sequencing of 64 populations from diverse climatic and agricultural zones further revealed substantial genomic variation associated with adaptation across environmental gradients and generally low genomic offset under future climate change, suggesting that the species is likely to persist as a yield-limiting agricultural weed under continued environmental change.

Niche tracking was the primary blackgrass response to climatic and anthropogenic changes in agroecosystems. This result aligns with previous reports that substantial niche evolution is rare among terrestrial plant invaders (Petitpierre et al., 2012; Wiens & Graham, 2005), and that 75% of plant taxa shifted their geographic distributions to track their preferred climates over the past 18 000 years (Wang et al., 2023). In biological invasions, introduced ranges frequently mirror native climatic niches, and niche evolution is typically modest (Early & Sax, 2014; Liu et al., 2020). For example, for *Ambrosia artemisiifolia*, populations introduced into Europe largely occupy niches similar to the native North American climatic niche, while Australian introductions occupy a subset of that niche, with little evidence for niche evolution (Putra et al., 2024). Moreover, environmental sorting of pre-adapted lineages can reproduce ancestry-climate clines across native and invaded ranges, indicating that broad climatic associations can remain conserved even as populations spread into new continents (Gamba et al., 2025).

In blackgrass, our models indicate that climate change increased the availability of suitable habitat for blackgrass, facilitating range expansion. Expansion was associated with environments exhibiting lower temperature and precipitation seasonality, higher precipitation in the wettest month, and higher mean temperature of the wettest quarter – conditions known to favour winter annual weeds such as blackgrass (Peters et al., 2014). The association with increased precipitation, reduced climatic variability and higher soil silt content suggests a preference for environments with greater water retention, consistent with blackgrass being more problematic in western and northern Europe (Moss, 2017). This is further supported by evidence that blackgrass outperforms wheat under waterlogged conditions (Harrison et al., 2024), as wheat is favoured on lighter soils (Stratonovitch et al., 2012).

Our genomic analyses provide insights into the mechanisms enabling persistence across environmental gradients. GEA revealed a polygenic basis for the species’ response to environmental variation, suggesting these populations could respond rapidly to and persist under mild, quick environmental shifts (Bellagio & Exposito-Alonso, 2025; Hayward & Sella, 2022; Kardos & Luikart, 2021; Barrett & Schluter, 2008; Gomulkiewicz & Holt, 1995). Candidate genes were enriched for diverse biological functions, including responses to abiotic and biotic stresses, likely facilitating persistence under variable and changing environments, including highly disturbed agricultural environments. Such adaptive variation may also serve as a reservoir of pre-adaptive alleles, enhancing the ability of blackgrass to respond to novel environmental challenges and future climate change (Cai et al., 2023; Dyer, 2018; Hawkins et al., 2019; Mohammad et al., 2022). This interpretation is supported by generally low genomic offset values.

Polygenic climatic adaptation is increasingly recognized across plant species experiencing spatial environmental heterogeneity and climate change. Similar patterns have been reported in the agricultural weed *Ambrosia artemisiifolia*, where thousands of genomic regions were associated with climatic variables across native and introduced ranges (Battlay et al., 2023); in *Lolium perenne*, where over ten thousand loci were identified as associated with 12 climatic variables (Blanco-Pastor et al., 2021); and in *Pinus ponderosa*, where more than 1000 loci showed association with five climatic variables (Shu & Moran, 2023). These studies suggest that polygenic climatic adaptation is widespread, although the genomic architecture underlying it may vary substantially among species.

The predominance of variable-specific associations suggests a largely modular genetic architecture, broadly consistent with expectations that heterogeneous selection tends to favour genetic modules responding independently to distinct environmental components (Le Nagard et al., 2011; Lotterhos et al., 2018). This combination of modularity and limited environmental pleiotropy may enable adaptation to complex environments, such as agroecosystems, by reducing maladaptive trade-offs among environmental responses.

Analyses combining species distribution modelling with genomic offset predictions revealed that most contemporary populations are genetically well aligned with future climatic conditions and suggested its likely persistence under future agroclimatic scenarios. Localized hotspots of elevated genomic offset occurred where projected climates diverged most strongly from historically suitable conditions, indicating some populations that may require greater genetic change to remain adapted. Similar spatial concordance between climatic novelty and genomic vulnerability has been reported in other species undergoing range shifts and environmental change (Capblancq et al., 2020), highlighting the value of integrating niche-based and genomic approaches to identify populations most vulnerable to future climate change. For example, in *Bromus tectorum* (cheatgrass), European native-range genotype-environment associations predict regions of high invasive dominance in western North America (Gamba et al., 2025). Our genomic offset results extend this to future climates, showing that most blackgrass populations remain well-matched to projected conditions, with only localized vulnerability.

Despite these strengths, several limitations warrant consideration. Historical occurrence records are uneven in sampling resolution and prone to taxonomic uncertainty, a well-recognized and inherent challenge in global biodiversity databases (Beck et al., 2014; Boakes et al., 2010; Sigler et al., 2021). While our curation and analytical design mitigate these issues, some uncertainty remains. Genomic offset metrics also rely on simplifying assumptions and do not fully capture the influence of demographic processes, migration, or adaptive plasticity (Gain et al., 2023; Láruson et al., 2022; Lotterhos, 2024; Rellstab et al., 2021). Integrating genomic time-series with experimental validation will help to refine predictions of long-term adaptive outcomes.

Taken together, our results highlight how ecological conservatism and evolutionary potential can jointly underpin species’ responses to environmental change in agroecosystems. Blackgrass exemplifies a scenario in which an agriculturally important species exhibits strong climatic niche conservatism across historical timescales, maintaining genomic variation to support adaptation across heterogeneous landscapes and facilitating range expansion under ongoing environmental change. This combination of ecological stability and evolutionary potential may be a characteristic of many successful agricultural weeds. More broadly, our integrative approach, combining historical niche reconstruction, ecological niche modelling, and landscape genomics provides a framework for understanding and forecasting adaptive responses to environmental change in agricultural systems.

## Acknowledgements

The authors thank everyone who collected and shared blackgrass seeds, namely Bayer Crop Sciences, Germany; David Comont, Rothamsted Research, UK; Husrev Mennan, Ondokuz Mayis University, Turkey; Katěrina Hamouzová, Czech University of Life Sciences, Czech Republic; Donato Loddo, Institute for Sustainable Plant Protection, National Research Council, Italy; Joel Torra Farré, Universitat de Lleida, Spain; Mette Sønderskov, Aarhus University, Denmark; Maor Matzrafi, Volcani Institute, Israel; and the United States Department of Agriculture, USA. We would also like to thank Guy Coleman for valuable discussions and comments. The project was supported by the Novo Nordisk Foundation (Grant number: NNF21OC0068600). The funders had no role in study design, data collection and analysis, decision to publish, or preparation of the manuscript.

## Data accessibility and benefit-sharing section

### Data accessibility statement

Genomic data generated *de novo* for this study are available on NCBI SRA under BioProject PRJNA1285609. GBIF data and code used in bioinformatic processing, analyses and data visualization is available in the GitHub repository https://github.com/celianeto/blackgrass_climate/.

### Benefit-sharing statement

Collaborations were established with scientists and institutions involved in sample collection, and their contributions are acknowledged in the manuscript. All genomic data and associated metadata have been made publicly available (see Data Accessibility Statement).

## Authors Contributions

CN designed the study, collected data, conducted all analyses and wrote the first draft of the manuscript. QZ conducted the wet-lab protocol. PN provided seed material and wrote the first draft of the manuscript.

