## Supplemental Information for "Climate-driven niche tracking and genomic resilience shape future distribution of a widespread agricultural weed"

#### Table of Contents:


| | |
|---|---|
| **Supplemental Methods** | Page 2 |
| **Supplemental Results** | Page 4 |
| **Supplemental References** | Page 7 |
| **Supplemental Figures** | Page 9 |

### Supplemental Methods

#### Genotype-environment association

Genome-environment association analyses were additionally performed using LFMM (latent factor mixed model) and RDA (redundancy analysis) as complementary methods to BayPass, to assess the robustness of candidate loci identification across methodological frameworks.

For both methods, SNPs were subsampled to one per 10kb genomic window, selected randomly, to reduce the influence of linkage disequilibrium on association tests. This reduced the number of variants considered from ~10M to ~183K. To account for the stochasticity introduced by this random subsampling and the considerably decrease in genomic coverage, all analyses were repeated across three independent replicates, each drawing a different random SNP per window.

To account for population structure, and since these two methods correct for it based on a fixed number of axes (principal components for RDA, latent factors for LFMM), we used the function *prcomp()* to calculate a population-level allele frequency matrix and then chose the best K based on the ‘elbow’ method. K = 4 was selected.

For LFMM, pool-seq allele frequencies were converted to pseudo-individual genotypes via beta resampling, as recommended for pool-seq data (LEA documentation, http://membres-timc.imag.fr/Olivier.Francois/lfmm/faq.htm), yielding a matrix of 3200 pseudo-individuals. Environmental variables were correspondingly replicated per population. LFMM2 was run separately for each environmental variable using the *lfmm2()* function from the LEA package (Caye et al., 2019; Frichot et al., 2013), and association p-values were obtained with *lfmm2.test()*.

For RDA, the same subsampled SNP sets as used in each LFMM replicate were analysed to ensure direct comparability. Population structure was accounted for using the same K as above. A separate pRDA model was fitted for each environmental variable using the *rda()* function from the vegan package (Oksanen et al., 2001), with neutral PCs included as covariates via the *Condition()* argument. SNP loadings on the single constrained axis were extracted with *scores()*, converted to z-scores, and transformed to two-tailed p-values assuming normality.

In both cases, p-values were corrected using the Benjamini-Hochberg FDR procedure, and SNPs with FDR < 0.05 were retained as candidates. Candidate SNPs were identified as those significant in at least one replicate in either method. To assess concordance with BayPass results, LFMM and RDA candidates were overlapped with BayPass significant genomic regions (expanded by ±10kb to account for linkage with causal variants) using positional intersection per environmental variable.

### Supplemental Results

#### Genotype-environment association analyses

Complementarily to BayPass, we conducted GEA analysis using LFMM and RDA approaches as well. Overall, LFMM identified 4380 SNPs in association with any variable (across the three replicates, average 1457 SNPs per replicate). No SNP was found in all three replicates (max 18 SNPs in common between any two replicates), showing how thinning can create false negatives depending on which SNPs are included in the analysis. RDA on the same set of SNPs across the three replicates identified 12 325 significantly associated SNPs. Both these analyses confirmed the strong polygenicity of climate adaptation seen with BayPass.

Overlapping these results – based on the same set of thinned SNPs – yield 16 SNPs identified across replicates and methods. Moreover, overlapping these with our initial BayPass results (and increasing the significant regions identified 10kb up and downstream) showed only one overlapping SNP (chr1: 723429722 for variable bio4). A two-way comparison showed 14 and 218 SNPs overlapping respectively for LFMM and RDA.

Both methods confirmed the polygenicity identified by BayPass but their overlap did not produce a *bona fide* set of validated candidates. Although triangulating GEA results is common practice, the observed lack of overlap is expected, as each method has its own assumptions, conditions and datasets. The differences between methods are not just mathematical but also biological, including differences in treatment of population structure, linkage disequilibrium, and allele frequencies in the population.

First – and most importantly – BayPass was explicitly designed to handle pool-seq data, implementing a binomial likelihood on read counts that natively accommodates sequencing depth uncertainty through the model (Gautier, 2015), a property absent from both RDA, which operates on point estimates of allele frequencies without modelling sampling variance, and LFMM, which is architecturally designed for individual genotype matrices (Caye et al., 2019; Frichot et al., 2013). Adapting LFMM to pool-seq data requires simulating pseudo-individuals via beta resampling of observed allele frequencies, which can introduce artificial within-pool variance and treats replicated draws from the same frequency estimate as independent observations, inflating the effective sample size seen by the latent factor model and potentially distorting both structure correction and association estimates.

Second, both RDA and LFMM correct for population structure by summarising the genotype matrix into a fixed number of linear axes (principal components for RDA, latent factors for LFMM) requiring the number of axes (K) to be specified in advance and assuming that population structure can be adequately captured by K discrete, linear components (Capblancq et al., 2018; Frichot et al., 2013). If K is misspecified, this can leave residual structure uncorrected or, conversely, absorb signal of interest into the structure-correction axes. BayPass instead corrects for population structure through a genome-wide covariance matrix Ω, estimated directly from pairwise allele-frequency covariances across all populations. This provides a continuous, non-parametric correction that captures complex demographic history (Gautier, 2015) without assuming a fixed number of discrete ancestral clusters or linear axes, and without requiring K to be chosen a priori.

Lastly, linkage disequilibrium poses a further challenge for GEA, as physically linked loci tend to show correlated association signals regardless of whether they are independently under selection. Standard implementations of RDA and LFMM address this through prior SNP thinning. This substantially reduces genome coverage, discards the majority of variants indiscriminately, and reduces resolution across entire genomic regions. Consistent with this expectation, three independent replicates of random thinning in our dataset showed no overlap in outlier SNPs even when using the same underlying method, demonstrating that thinning drastically reduces both coverage and power to detect associations. Moreover, in obligate outcrossing species such as blackgrass, where linkage disequilibrium is expected to decay rapidly, nearby SNPs may not be in strong linkage, meaning the retained representative SNP may not adequately represent the broader genomic window, risking the loss of genuine signal and a further reduction in power to detect polygenic associations. BayPass operates on all SNPs without prior thinning. Linkage disequilibrium was instead addressed using the Local Score approach (Bonhomme et al., 2019), which aggregates association signals across neighbouring loci while explicitly modelling their correlation structure, collapsing linked signals into discrete genomic regions without requiring variant removal. Since our primary goal is to characterise the full genetic architecture of climate adaptation –

including the extent and structure of environmental pleiotropy – maximising genomic coverage is essential. Restricting the analysis to a thinned subset would introduce ascertainment bias into overlap estimates between environmental variables, potentially underestimating pleiotropy where the causal variant happens not to be the retained representative SNP. Given these considerations, we retained BayPass results for all downstream analyses.

### Supplemental Figures

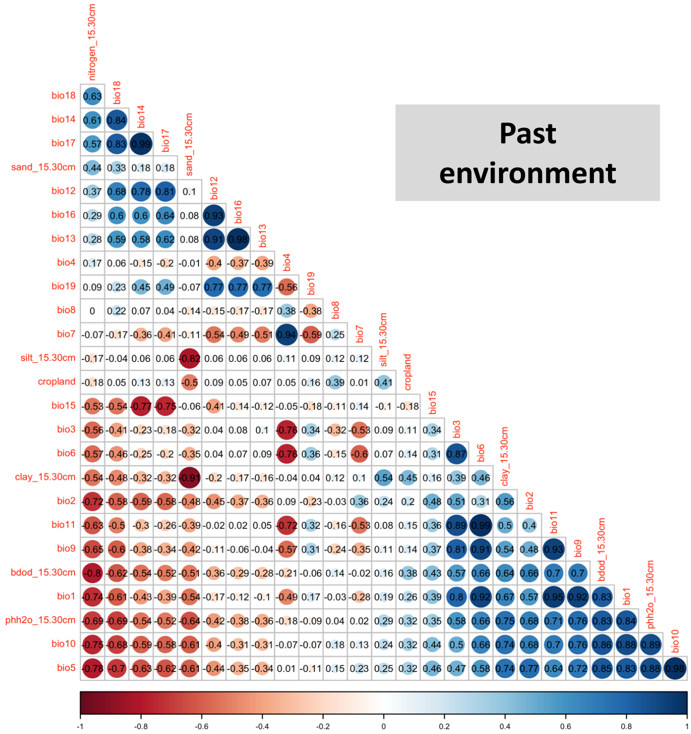


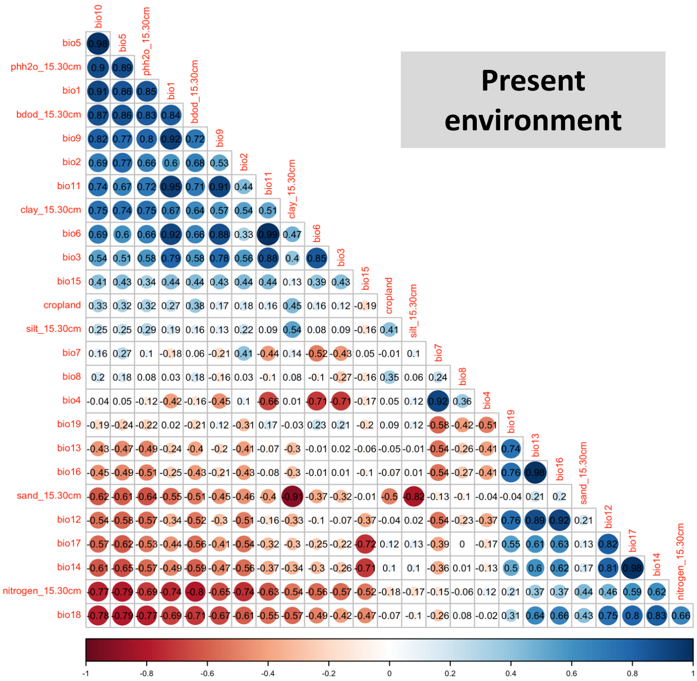


**Supplemental Figure 1. Correlation matrix between the 26 environmental variables considered in the past (top panel) and present (bottom panel).** Values show Spearman correlation.

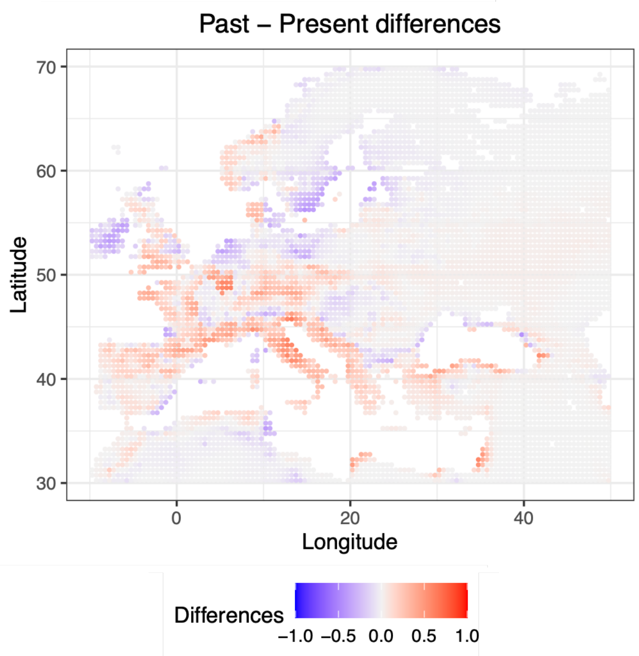


**Supplemental Figure 2. Differences in habitat suitability between the Past and the Present models.** Colour refers to differences in predicted suitability between the two models. A positive value represents higher suitability predicted under the Present model.

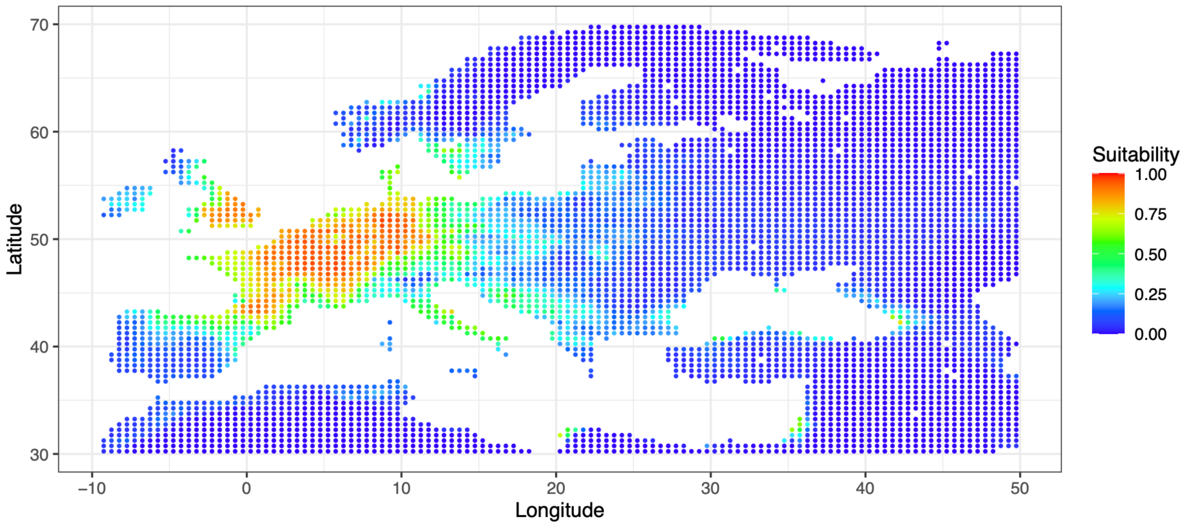


**Supplemental Figure 3. MaxEnt Future model.** Areas are coloured according to predicted suitability using present-day occurrences and future climate from the MPI-ESM1-2-LR, ssp 245 projection.

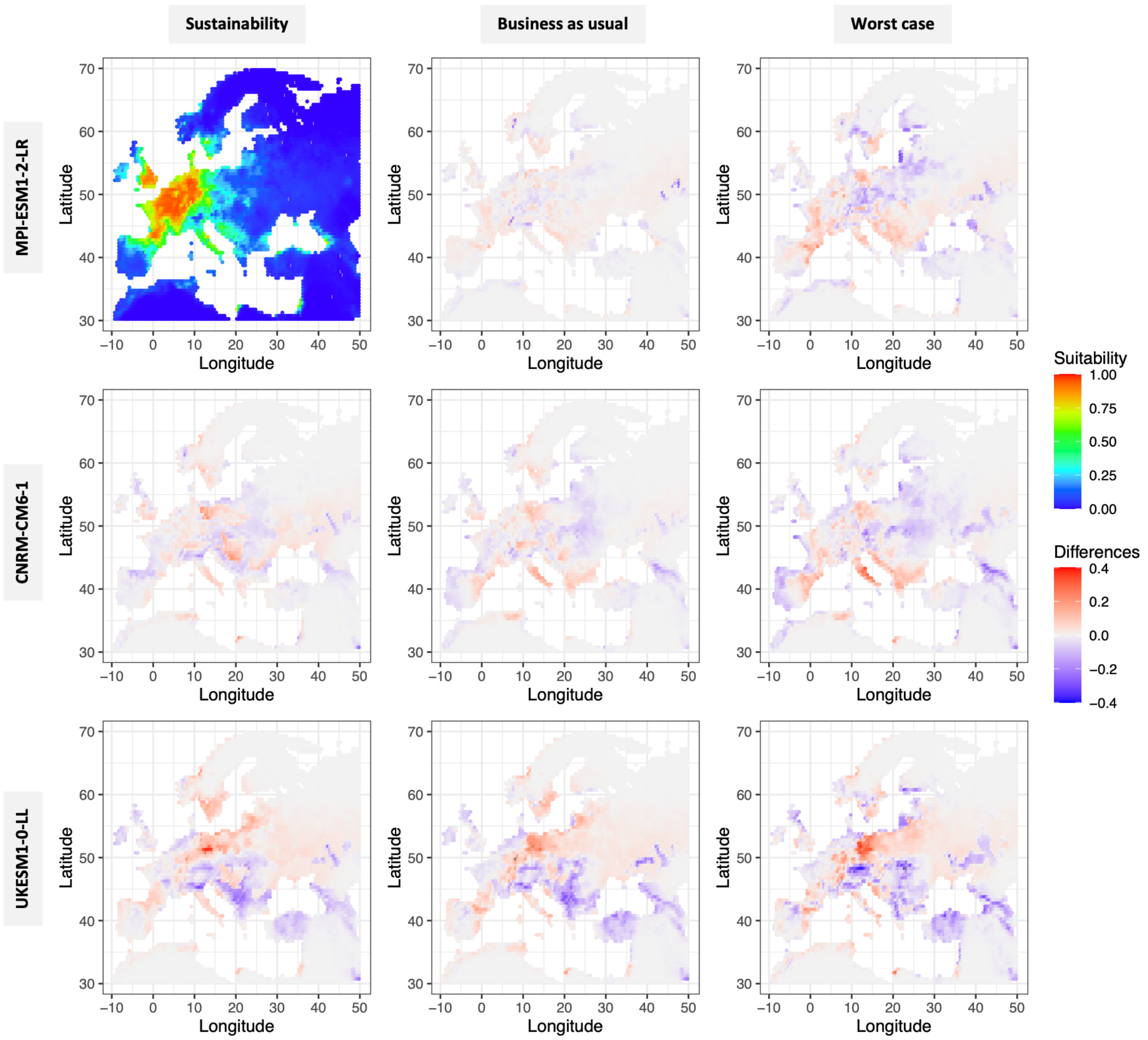


**Supplemental Figure 4. MaxEnt Future models for suitability**. Rows correspond to different models, according to the label on the left. Columns correspond to different emission scenarios (shared socio-economic pathways (ssp)), respectively, ssp = 126 (“Sustainability”), ssp = 245 (“Business as usual”), and ssp = 585 (“Worst case”). All models refer to the time period between 2061 and 2080. In each map, the colours refer to differences in suitability between each model and MPI-ESM1-2-LR, ssp 245 (the default model, top left corner). Red marks regions where suitability is higher for the default model, and blue regions where suitability is predicted to be lower.

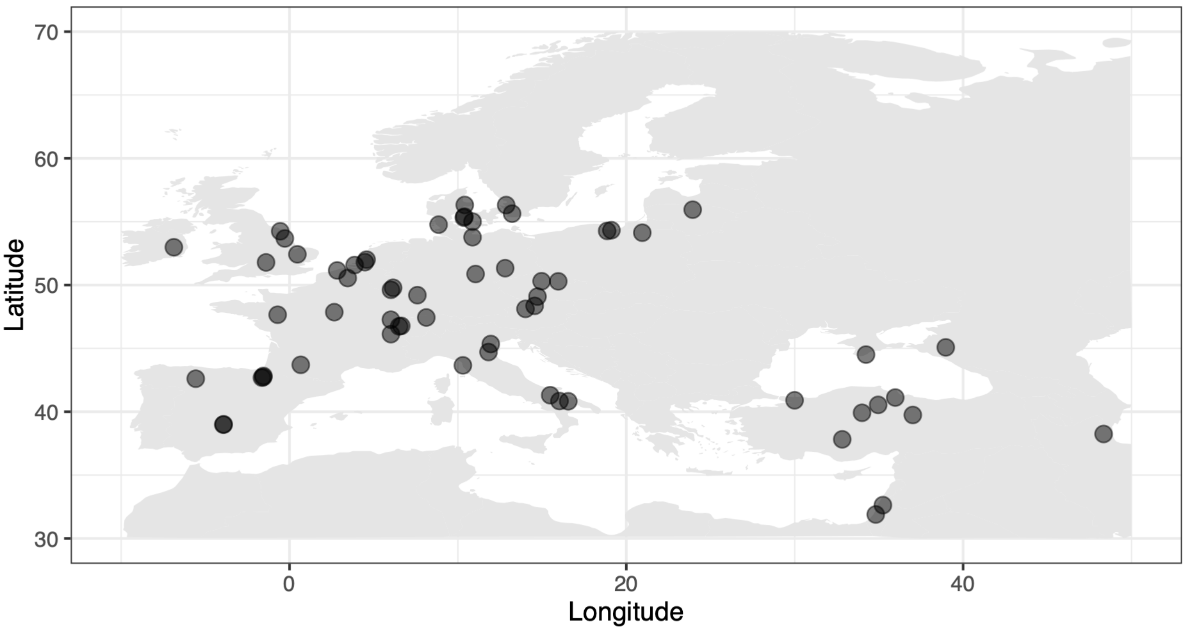


**Supplemental Figure 5. Geographical distribution of the 64 populations sampled and sequenced in this study.**

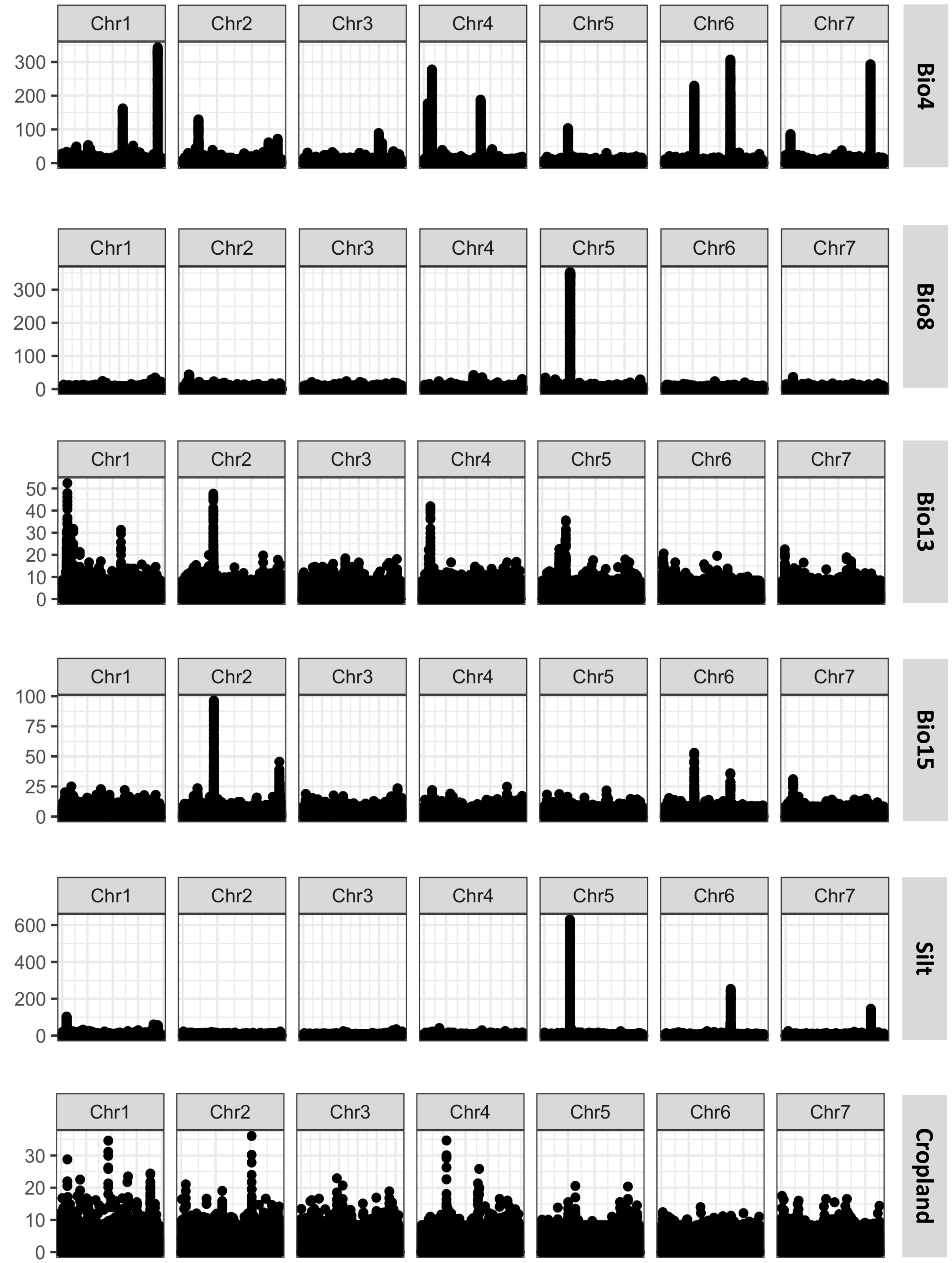


**Supplemental Figure 6. Manhattan plots for all mapped variables.** X-axis shows genomic positions and y-axis the Lindley score from the local score approach for each SNP. Labels with variable names are on the right: bio4: temperature seasonality; bio8: mean temperature of the wettest quarter;

bio13: precipitation of the wettest month; bio15: precipitation seasonality; silt: soil silt percentage, and cropland use.

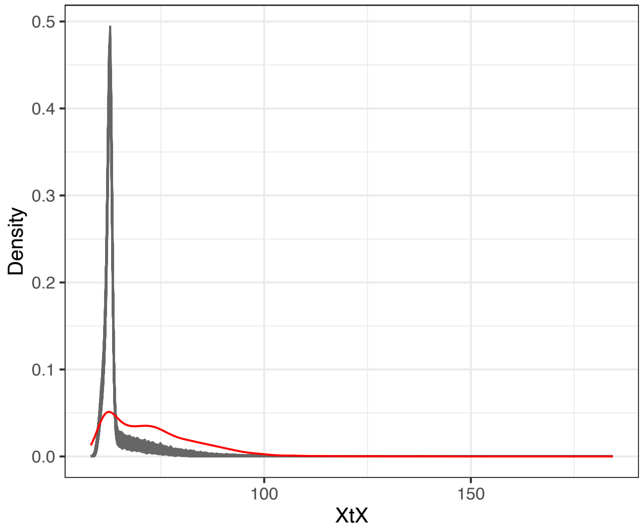


**Supplemental Figure 7. Environment-associated loci show signals of local adaptation.**

Distribution of XtX values (from BayPass) for each of the representative SNPs is shown by the red line. The grey distributions represent 1000 random draws from the genome-wide distribution of XtX values.

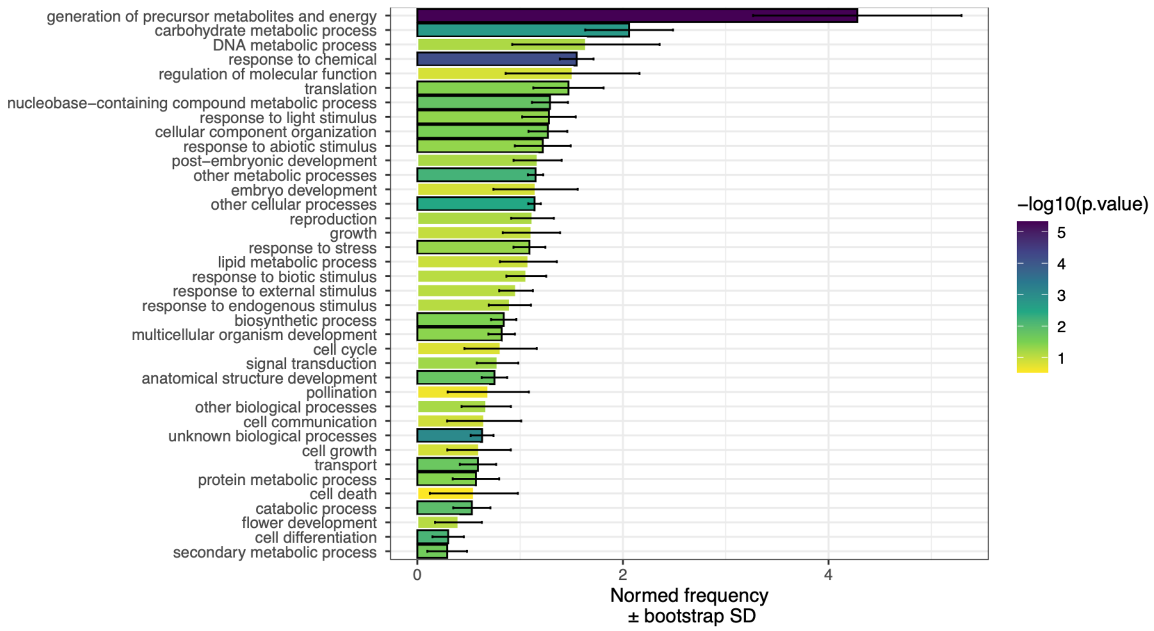


**Supplemental Figure 8. Gene ontology of candidate climate-associated loci.** Y-axis shows GO terms, while x-axis shows frequency with whiskers indicating bootstrapped standard deviation. Colours represent p-value, following the legend. Statistically significant GO terms are marked with a black contour on the corresponding bar.

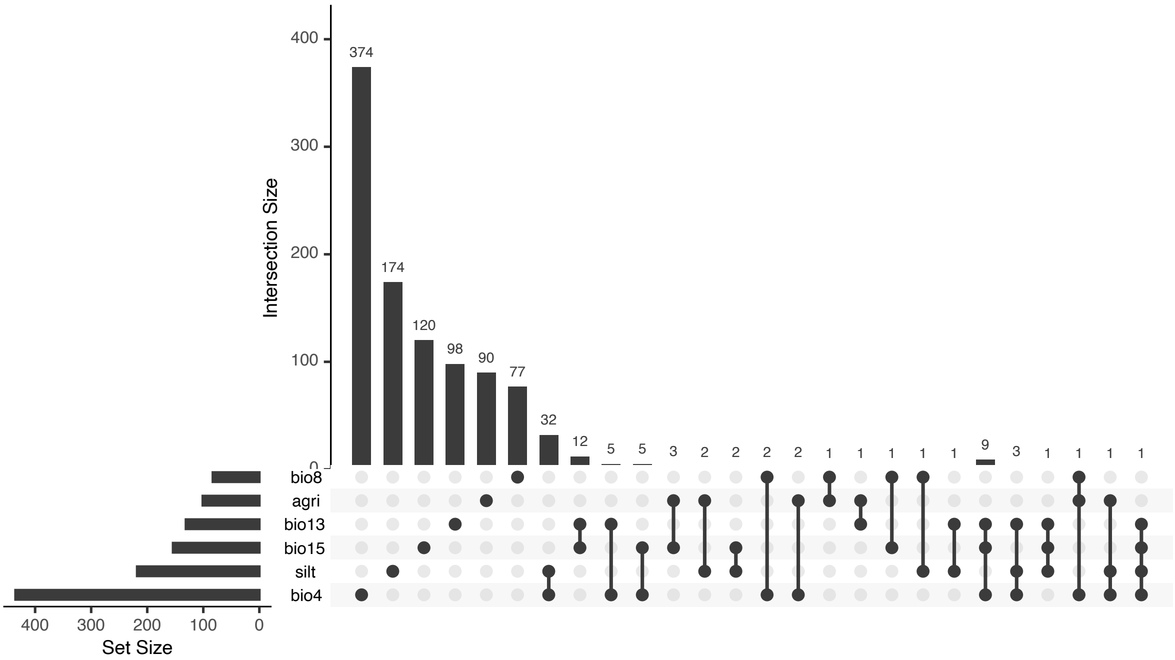


**Supplemental Figure 9. Environment-associated genes shared among variables.** Upset graph showing the number of different genes associated with each environmental variable (bar plot on the left) and their intersection across variables (bar plot on top). The dot matrix shows the number of genes shared between different variables. Environmental variables: bio4: temperature seasonality; bio8: mean temperature of the wettest quarter; bio13: precipitation of the wettest month; bio15: precipitation seasonality; agri: cropland use; silt: soil silt percentage.

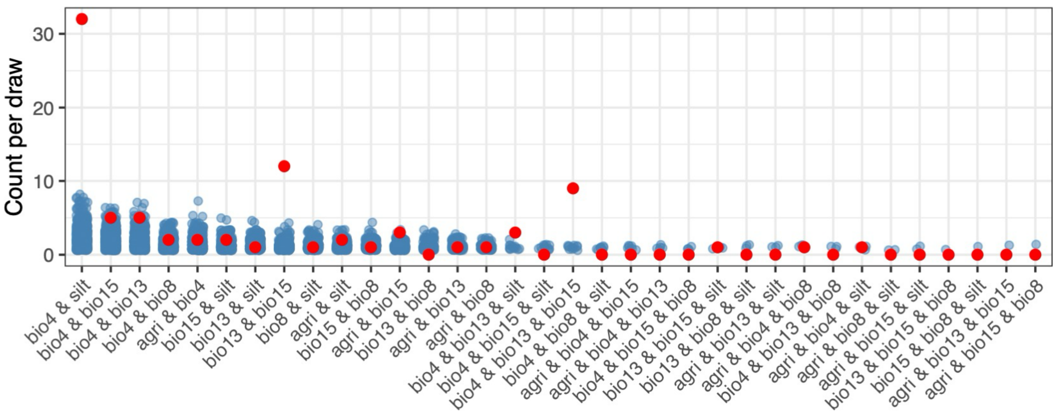


**Supplemental Figure 10. Null-model randomizations for multi-variable associations.** Counts of draws (in 1000, y-axis) for each combination of variables (x-axis) shown by the blue dots; the red dots represent the observed count of overlap between variables from GEA results.

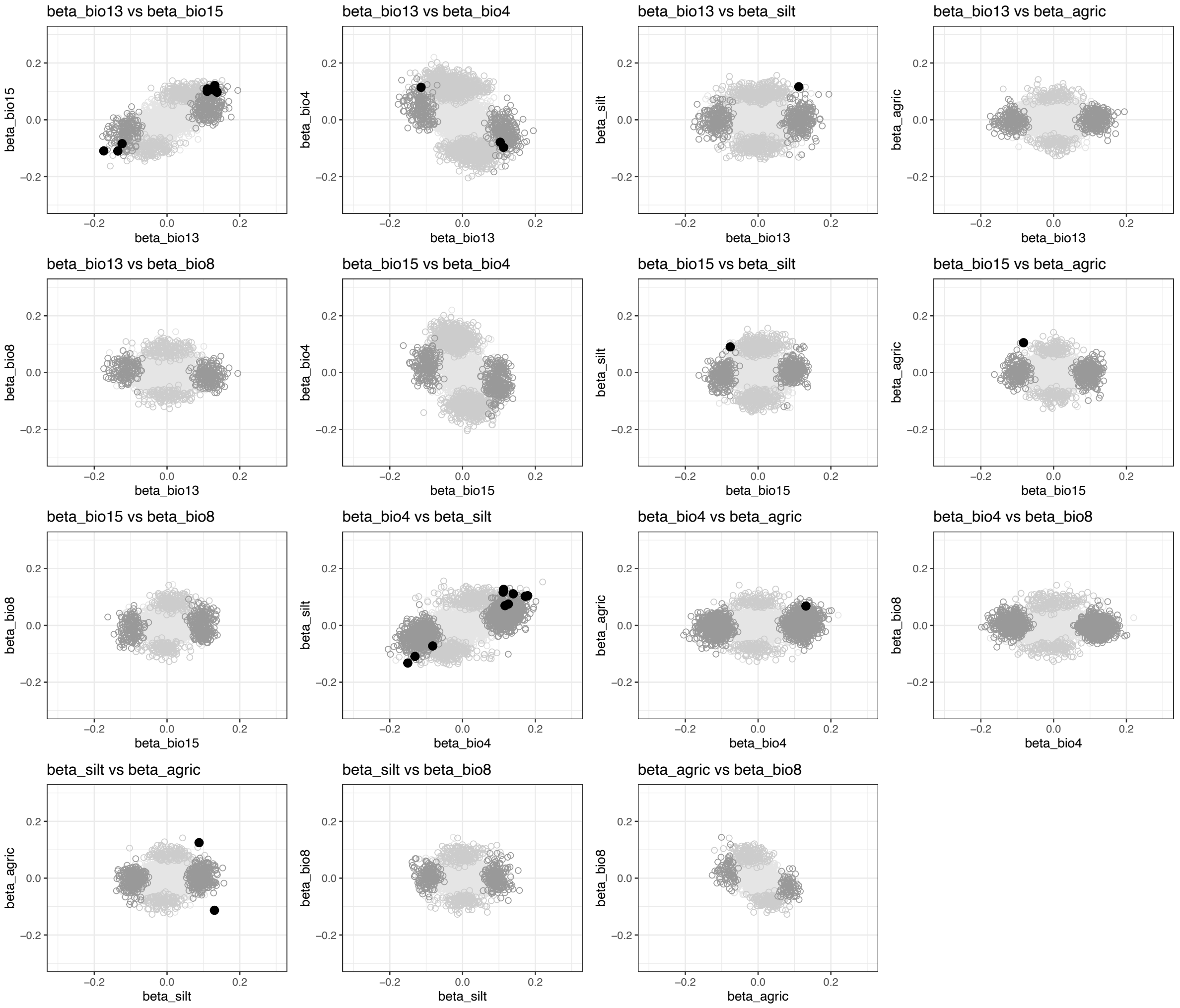


**Supplemental Figure 11. Genetic correlation between environmental variables for the 3523 representative SNPs.** Each panel shows genetic correlation between any two environmental variables. For each SNP, represented by dots, effect size (beta, calculated in BayPass) is shown for the two environmental variables considered per panel (variables names shown on the axes and panel title). Lightest grey colours mark SNPs not correlated with any of the two considered variables per panel. The intermediate shades of grey show SNPs associated with one but not with the other variable. Black dots represent SNPs with statistical association with the two considered variables. Environmental variables: bio4: temperature seasonality; bio8: mean temperature of the wettest quarter; bio13: precipitation of the wettest month; bio15: precipitation seasonality; agri: cropland use; silt: soil silt percentage.

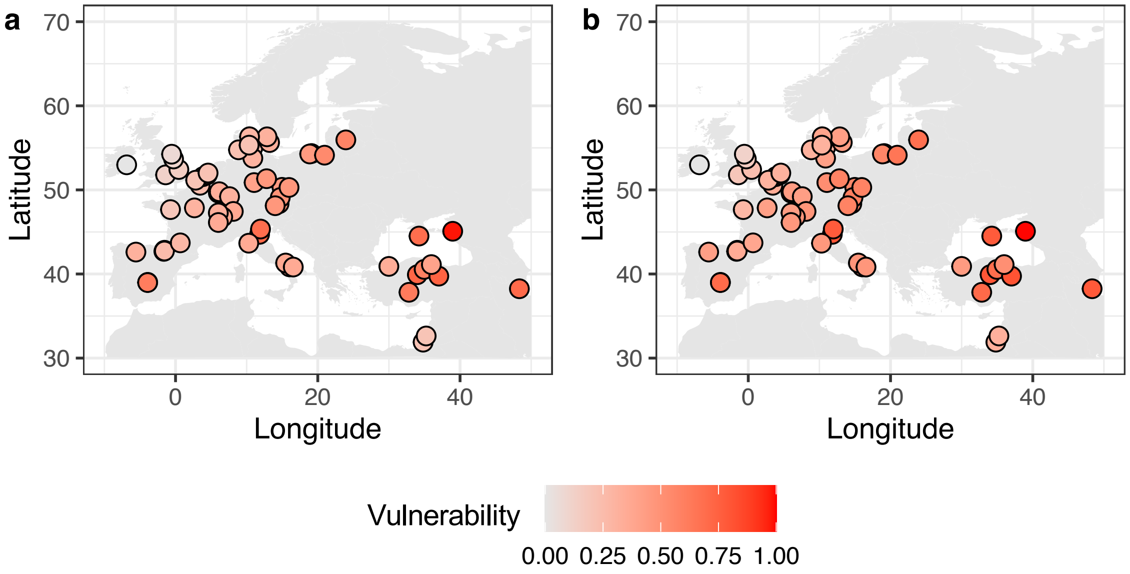


**Supplemental Figure 12. Comparison between methods to estimate genomic offset. a.** Geometric genomic offset, and b. RONA (risk of non-adaptedness). Each dot marks one population from the 64 included in this study, coloured by the corresponding genomic offset metrics. Red shows populations with higher estimated genomic offset, and therefore more vulnerable to climate change. Values were scaled to allow for comparisons across approaches.

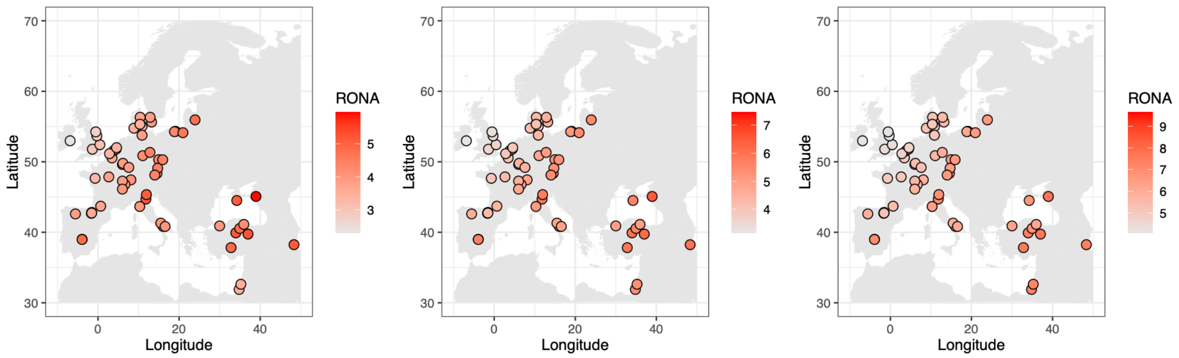


**Supplemental Figure 13. Risk of non-adaptedness (RONA) across the 64 populations under different XtX-based candidate SNP thresholds.** From left to right, the panels show all candidate climate-associated SNPs included in the genomic offset model (left), the top 10% of SNPs by XtX value (middle), and the top 1% of SNPs by XtX value (right).

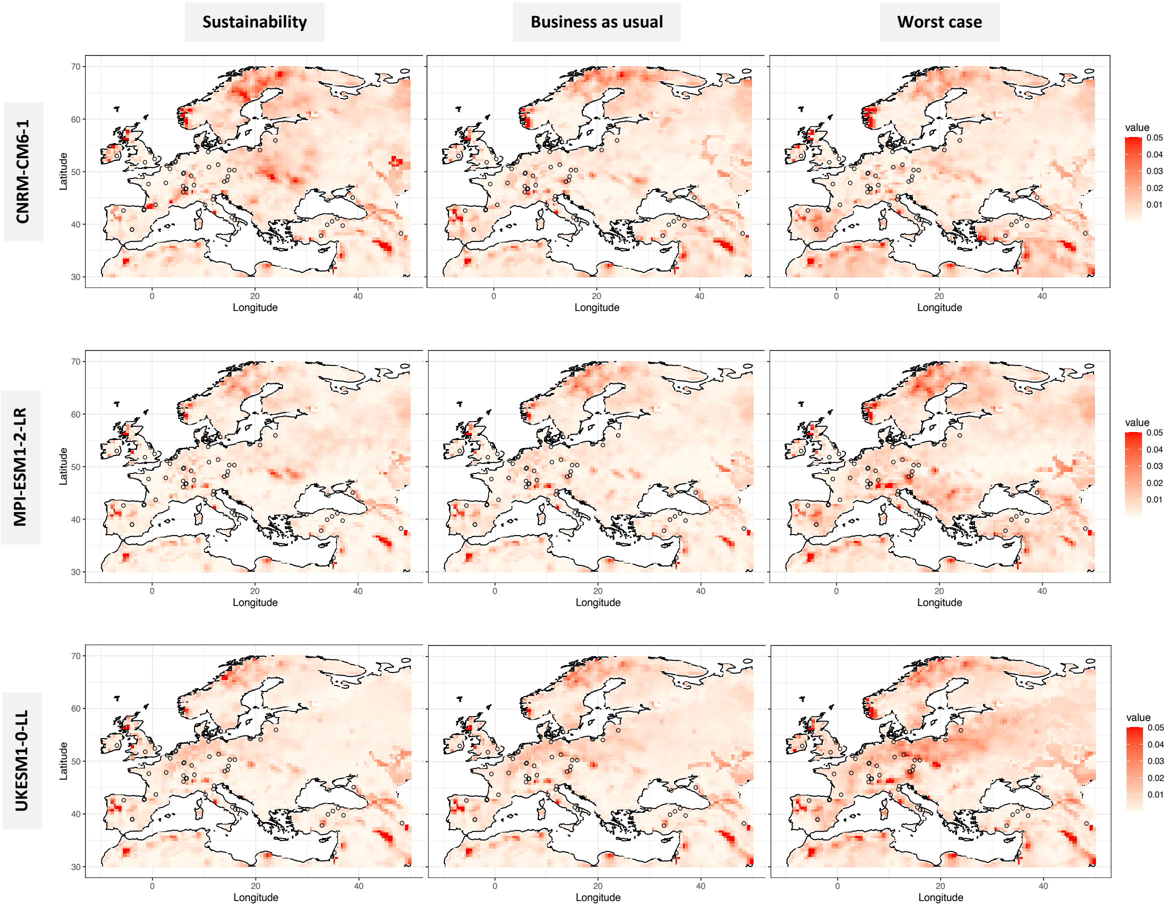


**Supplemental Figure 14. Genomic offset in Europe under different future climate models.** Rows correspond to different models, according to the label on the left. Columns correspond to different emission scenarios (shared socio-economic pathways (ssp)), respectively, ssp = 126 (“Sustainability”), ssp = 245 (“Business as usual”), and ssp = 585 (“Worst case”). All models refer to the time period between 2061 and 2080. In each map, the circles show the 64 natural populations used in the analysis. Shades of red represent genomic offset values computed with the geometric genomic offset approach.
